# Levodopa and dopamine dynamics in Parkinson’s disease metabolomics

**DOI:** 10.1101/306266

**Authors:** Rachel C Branco, William Ellsworth, Megan M Niedzwiecki, Laura M Butkovich, Douglas I Walker, Daniel E Huddleston, Dean P Jones, Gary W Miller

**Affiliations:** Department of Environmental Health, Rollins School of Public Health, Emory University, Atlanta, GA, 30322, United States; Department of Physiology, Emory University School of Medicine, Atlanta, GA, 30322, United States; Clinical Biomarkers Laboratory, Division of Pulmonary, Allergy, Critical Care, and Sleep Medicine, Emory University, Atlanta, GA, 30322, United States; Department of Neurology, Emory University School of Medicine, Atlanta, GA, 30322, United States; Department of Environmental Health, Rollins School of Public Health, Department of Pharmacology, Department of Neurology, Center for Neurodegenerative Diseases, Emory University, Atlanta, GA, 30322, United States

**Keywords:** Parkinson’s disease, metabolomics, levodopa, dopamine

## Abstract

Parkinson’s disease (PD) is a progressive neurological disorder caused by a combination of genetic and environmental factors. Metabolomics is a powerful tool that can be used to screen for potential biomarkers, exogenous toxicants, and metabolic network changes associated with disease states. Here, we used high-resolution metabolomics to compare over 10,000 plasma metabolic features from older adults with and without PD in an untargeted approach. We performed a network analysis that demonstrates that the presence of the PD drug levodopa influences variation observed between PD and control patients. Metabolome wide association studies and discrimination analysis identified significant differentiation in the metabolomics profile of older adults with and without PD. Notably, 15 metabolic features (ten of which we putatively identified) differed between PD and control adults with *p* < 0.05 and a corrected false discovery rate less than 20%. Furthermore, 13 metabolic networks were identified to be functionally different between PD and non-PD patients. Lastly, the dopaminergic toxic intermediate DOPAL differed between PD and non-PD populations, which supports the dopaminergic sequestration model of PD. These individual metabolites and metabolic networks have been implicated in past PD pathogenesis models, including the beta-carboline harmalol and the glycosphingolipid metabolism pathway including the ganglioside GM2. We recommend that future studies take into account the confounding effects of levodopa in metabolomic analyses of disease versus control patients, and encourage validation of several promising metabolic markers of PD.

---

Parkinson’s disease (PD) is a common progressive neurodegenerative disorder. Though PD pathology affects multiple systems, the most distinctive symptom of PD is motor dysfunction, including bradykinesia, rigidity, and postural instability [1]. PD is caused by the loss of nigrostriatal dopamine neurons [2], though other brain regions are affected in PD pathology [3,4]. At the time of PD diagnosis, approximately 60% of nigrostriatal dopamine neurons are already lost [5]. To counteract the loss of dopamine neurotransmission after dopamine cell loss, patients take an oral medication containing levodopa (L-dopa), a dopamine precursor which can cross the blood-brain barrier and helps overcome the paucity of endogenous dopamine in the brain [6]. In this way, L-dopa acts to slow the progression of motor symptoms [7]. However, there are no treatments to either halt or reverse PD progression. Despite the cluster of unwanted complications associated with treatment, L-dopa remains the mainstay pharmacotherapy for PD.

Because earlier identification of PD is critical to effective treatments, research has focused on the identification of biomarkers, protective factors, and/or relevant pathways that characterize PD risk and pathology. High-resolution metabolomics methods provides a powerful tool to characterize the molecular profile present in biological samples. Untargeted metabolomics can simultaneously characterize thousands of endogenous and exogenous compounds within a biological sample – collectively called the *metabolome* [8–11].

Due to the treatment considerations of sampling human patients, most metabolomics studies of human PD have sampled from PD patients that are actively taking L-dopa or that are L-dopa deprived for a only short period of time prior to sample collection [12–14]. The first aim of this study was to test how much L-dopa accounts for the metabolic differences between PD patients and controls. We found that the majority of metabolic profile differences between Parkinson’s and control patients can be explained by the presence of L-dopa, even in metabolic networks theoretically independent from dopamine metabolism. In this way, this paper acts as a warning to future biomarker and metabolomics researchers to account for the effects of treatment when evaluating effects of pathology.

The second aim of this study was to evaluate biomarker metabolites or pathways that differ between PD and control patients. Despite the complicating effects of L-dopa, we identified metabolic differences that are likely due to PD pathology rather than the L-dopa treatment. Our findings are consistent with past experimental literature suggesting that the antioxidant beta-carboline harmalol and the glycosphingolipid pathway are altered in PD and provides the first demonstration showing these metabolic variations are detectable in peripheral blood. Furthermore, this metabolic analysis underscores the altered dopamine dynamics present in PD patients.

Author summary: Prognostic diagnosis of disease is key for identifying treatment options that slow disease progression and improve patient quality of life. This is particularly important in Parkinson’s disease, where patients could greatly benefit from treatment before their latent sickness becomes problematic. Past studies have compared the blood of Parkinson’s patients with the blood of healthy adults in an effort to find a biomarker for Parkinson’s disease—however, in most of these past studies, the patients with Parkinson’s disease were taking the Parkinson’s medication levodopa. In our study, we analyzed the plasma metabolomics profiles of people with and without Parkinson’s disease. We found that the presence or absence of levodopa was the main difference between the blood of people with and without Parkinson’s disease. Furthermore, levodopa was associated with alterations in other metabolic pathways in a way that makes it hard to determine which differences are due to the disease and which are drug-related. Despite this complicating factor, we identified compounds and pathways that differ between Parkinson’s and control patients, including harmalol and the glycosphingolipid pathway. With further testing, these markers may help doctors identify Parkinson’s risk earlier in life.

## Results

There were no differences in the proportions of men and women between the PD and control group, but differences were found in age, race, and educational level, with the PD group more likely to be Caucasian, younger, and less educated. As expected, PD and control groups also differed in motor and non-motor symptoms of PD (Table 1).

**Table 1.**
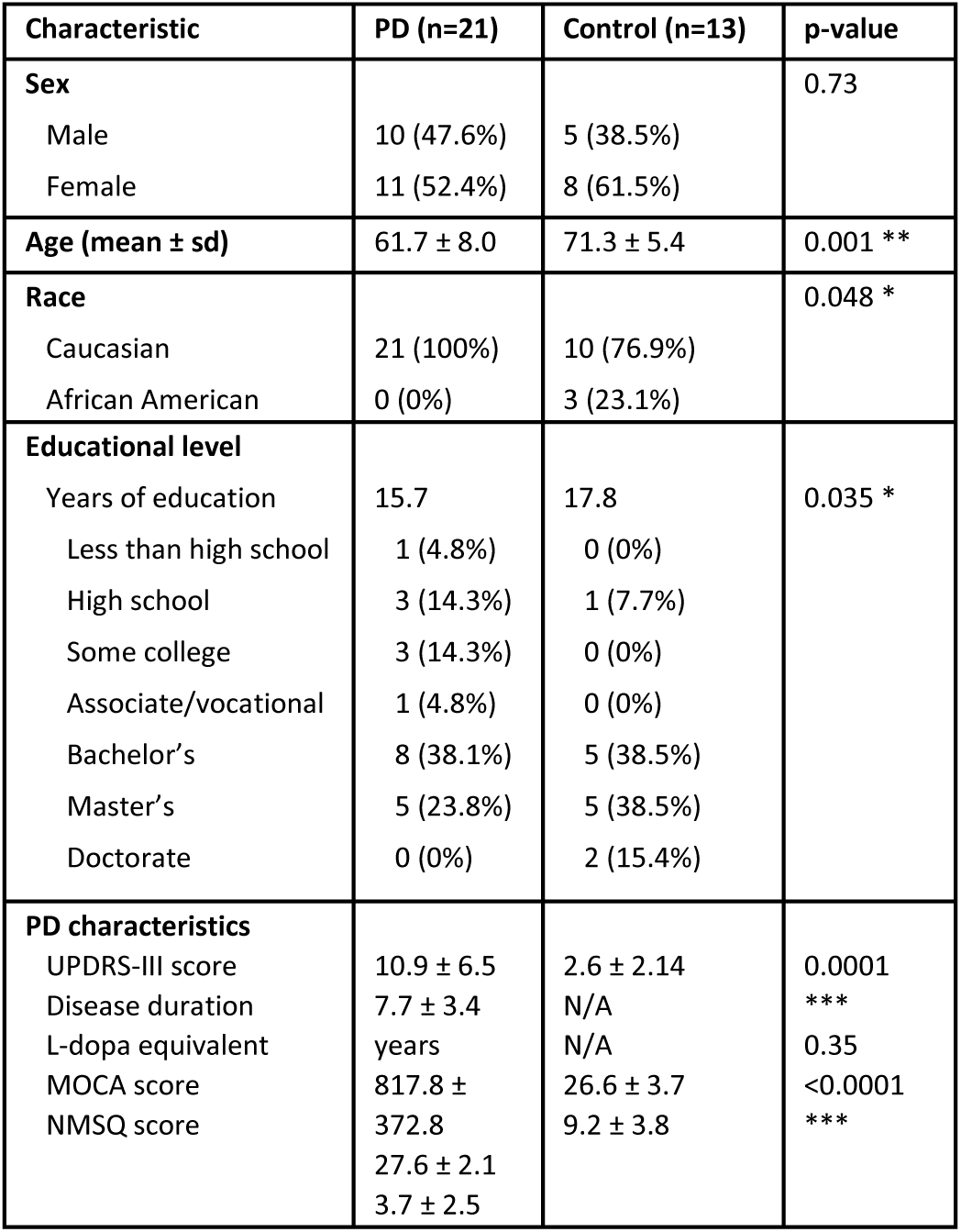
Description of study participants. 21 PD patients and 13 aged controls were studied. Differences between groups were calculated by Fisher’s exact test for Sex and Race, and t-test for all other categories.

We performed t-tests to identify *m/z* features differentially expressed in patients with PD based upon a false discovery rate (FDR) correction threshold of 20%. Fifteen *m/z* features were different between the two groups (Fig 1, Table 2). Predictably, the five top annotated compounds were either PD drugs or their metabolites, all of which were elevated in PD patients (Fig 2A). These PD drug metabolites met at least 2 criteria for identity, i.e., accurate mass match to predicted adduct (Level 5 identity [15]), co-elution with authentic standard, ion dissociation spectrum matching that of known compound, or association with known pathway or metabolic network (Supplementary Table 1), are indicated by metabolite name. Other features are denoted by associated accurate mass match to known metabolite (Level 5 identity) along with *m/z* and retention time, as indicated. Additionally, the toxic dopamine metabolite 3,4-dihydroxyphenylacetaldehyde (DOPAL) was elevated in PD patients (Fig 2B). Several compounds, including harmalol, were elevated in control patients relative to PD patients (Fig 2C). Five of the fifteen metabolic features provided no matches in chemical databases.

**Table 2.**
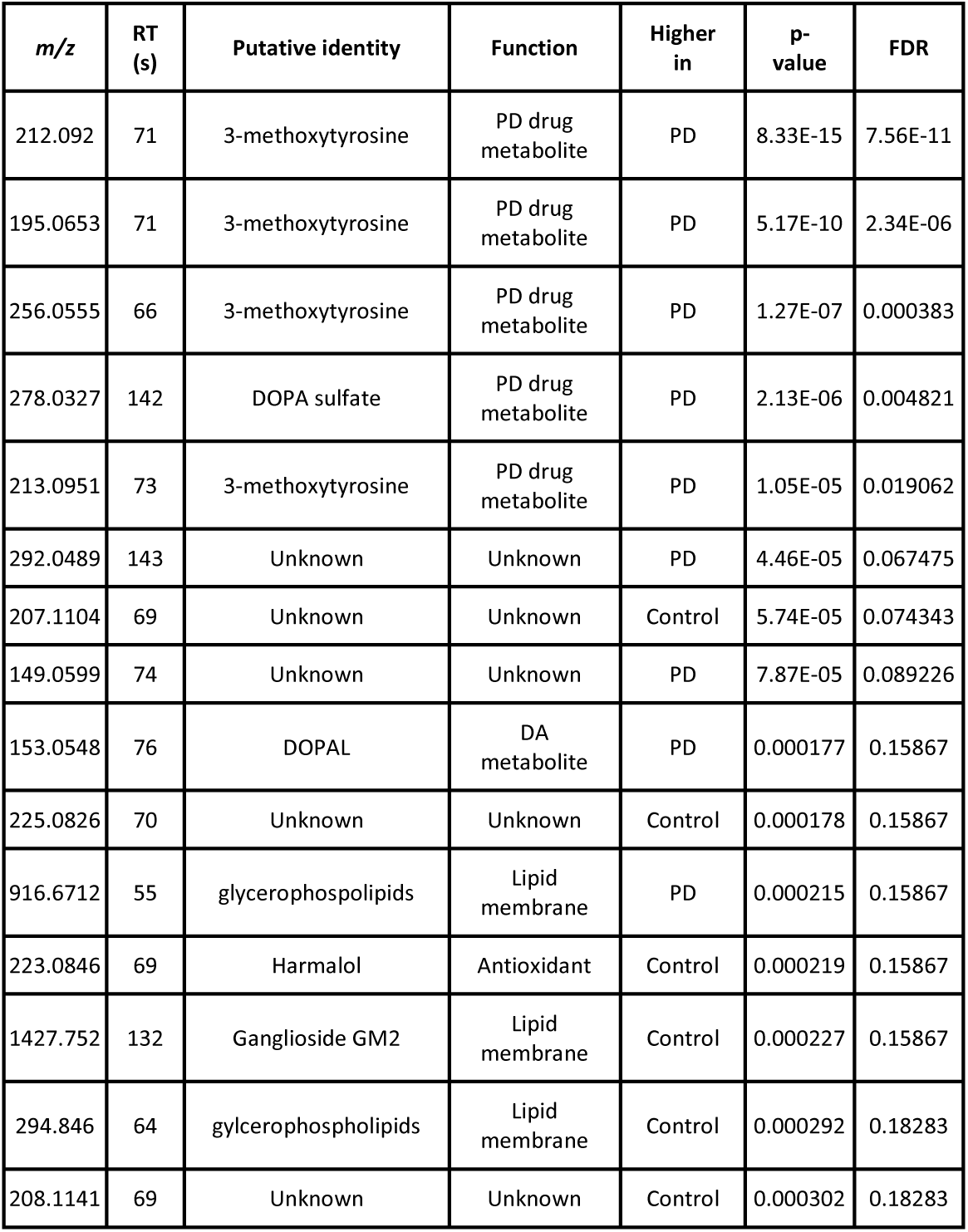
15 compounds resulted from the untargeted MWAS comparing features in PD patients against control patients. Using the Human Metabolome Database (HMDB), likely candidates for 10 of the 15 metabolic features were identified; the remaining 5 were provided no matches.

**Fig 1.**
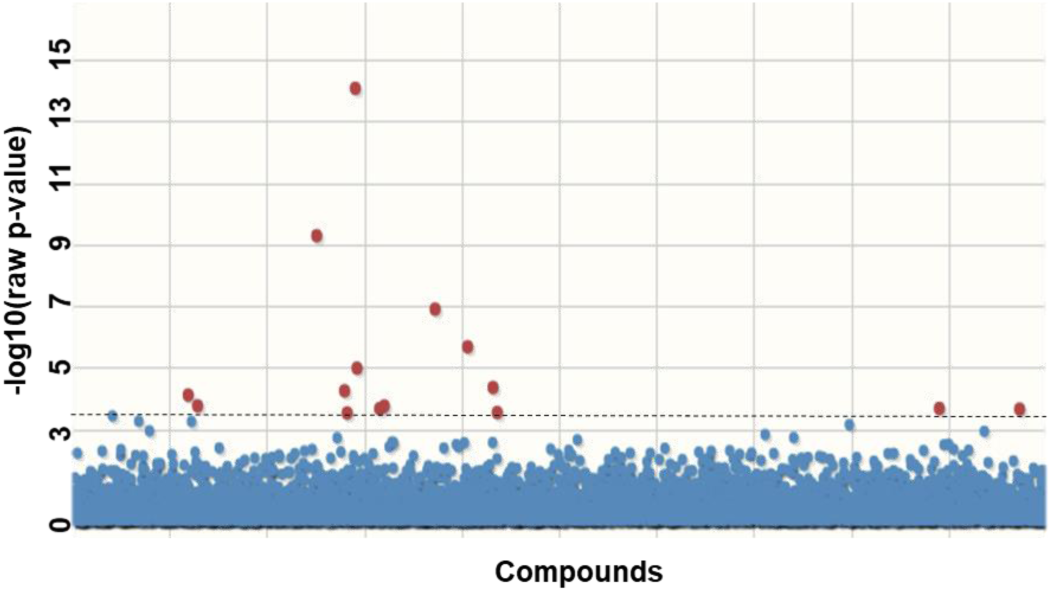
Manhattan plot for the untargeted MWAS. The x axis contains features arranged in order of *m/z*, and the y axis contains −log10 of the unadjusted p value.

**Fig 2.**
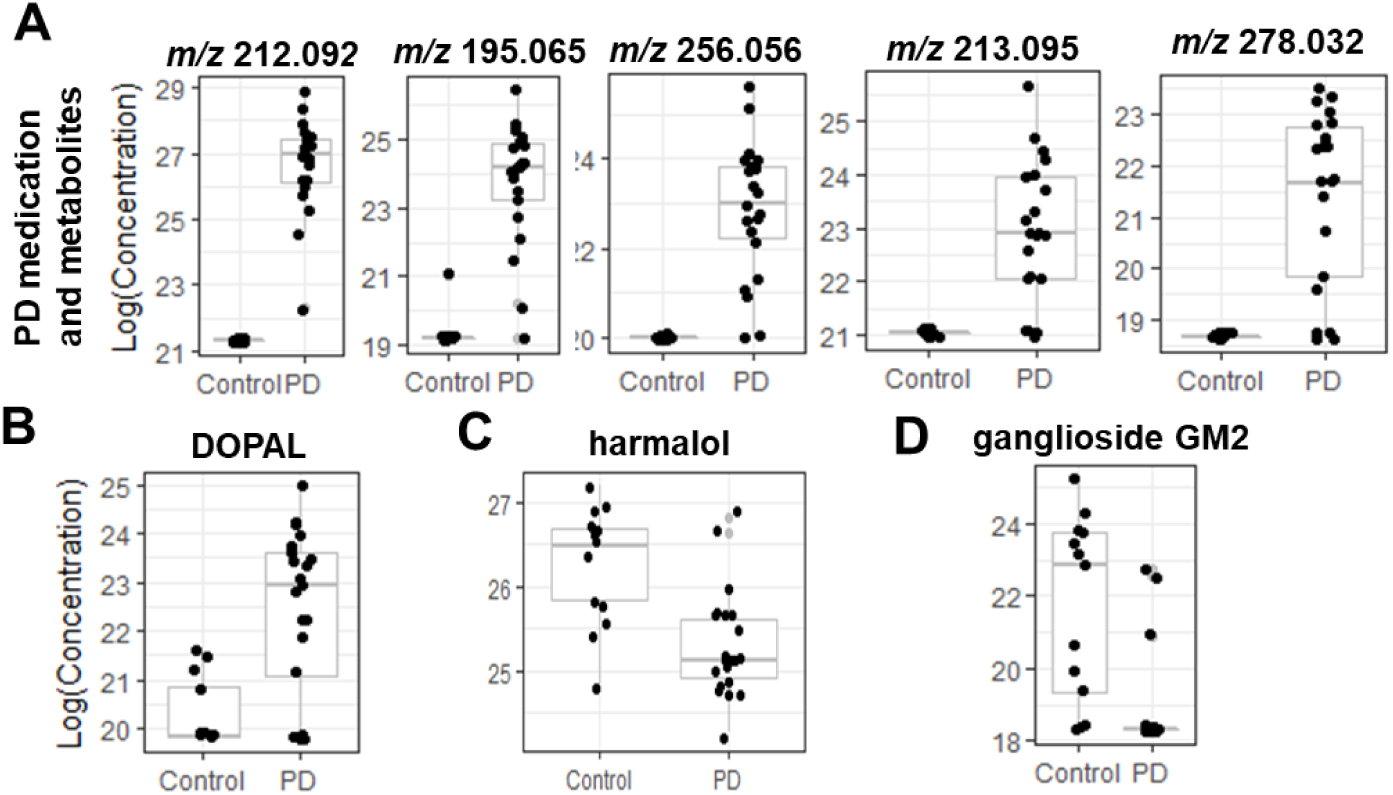
Boxplots for the 8 annotated compounds from the untargeted MWAS. The x axis contains clinical status and the y axis contains Log10(Concentration).

Each patient’s L-dopa equivalent dose was highly correlated with the primary L-dopa metabolite (*m/z* 212.092, r^2^ = 0.6559). Furthermore, L-dopa metabolite abundance and Unified Parkinson’s Disease Rating Scale Part III (UPDRS) scores were highly correlated (r^2^ = 0.4562), indicating that, as expected, patients with more severe PD symptoms tended to have more L-dopa in their blood.

We performed orthogonal partial least squares-discriminate analysis (OPLS-DA), which uses supervised learning to determine the principal components that most differentiate populations across multiple dimensions [16]. Using OPLS-DA, the metabolic profile of PD and control patients differentiated into two distinct groups (Fig 3). Within the OPLS-DA analysis, we identified features that most contributed to the model using the covariance and correlation between the features and the class designation. As expected, the features that were most influential in driving the OPLS-DA model corresponded to the features identified as most significantly different between PD and controls in the t-test analysis. To evaluate if OPLS-DA merely stratified patients according to presence of L-dopa, we completed a second OPLS-DA after manually eliminating signals corresponding to L-dopa or direct L-dopa metabolites. Even with these PD drug features removed, the two populations differentiated into two distinct groups (Supplementary Fig 1).

**Fig 3.**
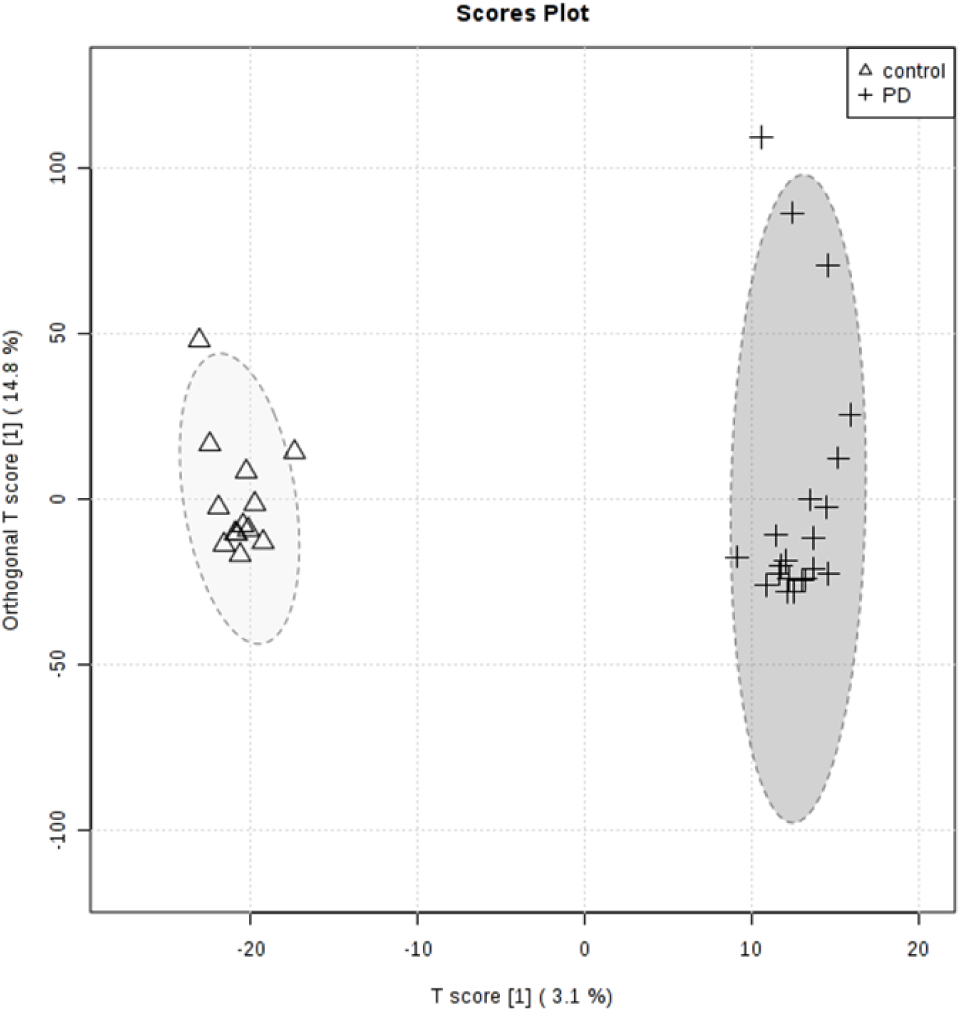
Orthogonal Partial Least Squares Discriminant Analysis (OPLS-DA). Clear separation between the PD and Control group were visible.

However, the removal of L-dopa and its direct metabolites from the OPLS-DA analysis does not guarantee that the downstream effects of L-dopa are eliminated. In order to test the correlation between L-dopa and other metabolites, we performed a separate analysis to predict which metabolites correlated with L-dopa (Supplementary Fig 2). We then performed pathway analysis to elucidate the metabolic pathways that were altered between PD and control patients (Supplementary Fig 3). By comparing the results of these metabolic pathway analyses, we identified several major metabolic pathways altered in PD patients and correlated with plasma L-dopa (Fig 4, Supplementary Table 2). Notably, every metabolic network that was different between PD and control plasma metabolites was also affected by the presence of L-dopa.

**Fig 4.**
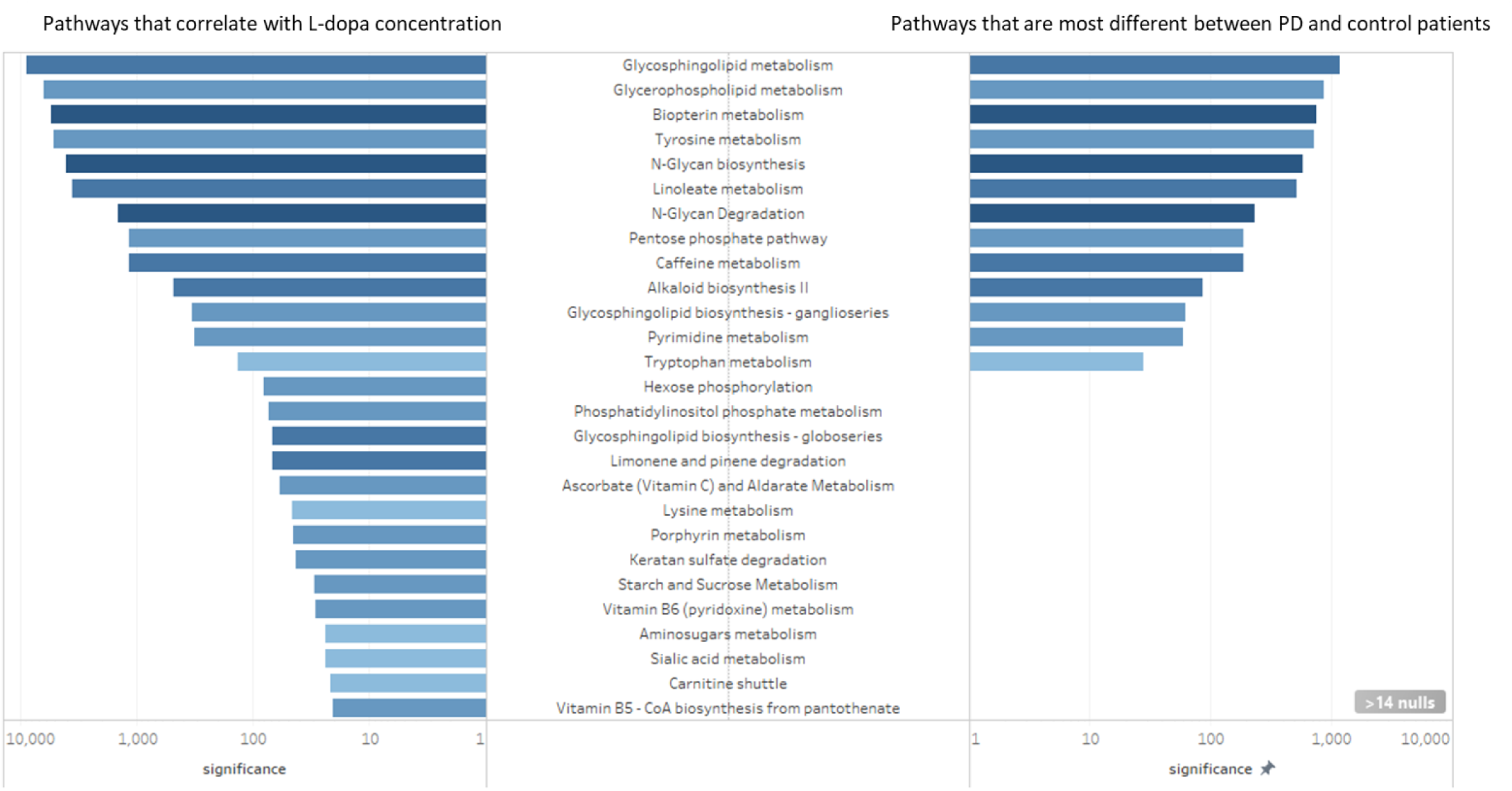
Altered metabolic pathways. Left, metabolic pathways that correlate with L-dopa concentration. Right, metabolic pathways that are most different between PD and control patients. The x-axis is degree of significance, calculated by log(1/adjusted-p-value). Metabolic pathways with adjusted p-values greater than 0.05 were excluded from visualization. Color corresponds with the extent that the network was affected, where dark blue denotes that a large proportion of metabolites within the network were affected, and a light blue denotes that a relatively small proportion of metabolites within a network were affected.

## Discussion

### L-dopa confounds PD versus control comparisons

The top five differential features measured in PD and control patients’ plasma were a PD drug and its metabolites (Table 2). The elevated levels of these L-dopa metabolites in PD patient plasma acted as a de facto positive control. The known oral dose of L-dopa strongly correlated with *m/z* 212.092, which was identified as 3-methyltyrosine. Therefore, we deemed the *m/z* 212.092 feature as an acceptable proxy for L-dopa concentration in patient plasma samples:

Feature reduction analysis using OPLS-DA showed that the metabolic profiles of PD versus control patients were distinctly different across two principal components (Fig 3). We also performed OPLS-DA on PD versus control patient plasma metabolites after manually removing the metabolic features corresponding to L-dopa or direct L-dopa metabolites. This approach also resulted in distinct clustering of PD and control populations across two principal components. However, this result does not prove that PD drugs did not drive the separation of the two groups, as the biological response to L-dopa could have driven the separation between PD and control groups despite the omission of L-dopa itself. Indeed, L-dopa is correlated with many metabolic networks, so it is likely that downstream effects of L-dopa alter critical metabolite features that define each metabolic profile and underlie the OPLS-DA identified differences between PD and control patients.

The comparison of metabolic pathways between PD and control patients revealed a number of biological pathways that were different in PD. However, in this analysis, it was impossible to tell whether metabolic pathways are altered due to PD pathology or by PD treatment. To this end, we performed a separate pathway analysis restricted to PD patients to identify which biological pathways were most strongly associated with L-dopa. All pathways that were identified using metabolites that were different between PD and control patients were also correlated with the presence of L-dopa. This includes networks that we would expect to be perturbed by L-dopa side administration, such as tyrosine and biopterin metabolism. However, many of these pathways were correlated with L-dopa despite the lack of any *a priori* suspicion of biological interaction. In previous studies, these pathways were assumed to be altered in PD patients due to PD pathology, but the analysis shows that L-dopa is a confounding factor to that interpretation.

These results provide a warning to future metabolomics studies using prevalent disease cases the extent to which treatment of the disease confounds identification of underlying pathology. Since L-DOPA levels correlate well with many non-dopamine metabolic pathways, simply removing L-DOPA from the dataset after the data collection is likely not sufficient to reduce confounding effects of the drug.

It is not surprising that the treatment of the condition (in this case, L-dopa for PD) has a close relationship with pathways that are pathological in the disease state (i.e. dopamine metabolism). Dopamine dynamics are undoubtedly altered in PD patients [2,3,17] — it is impossible to tease apart the contribution of the treatment and of the disease in this paradigm. A small number of studies have looked at metabolomics of peripheral fluids in PD [12–14], few of which include PD patients without L-dopa [18–20]. Several studies have looked at plasma or urine metabolites in PD versus controls, and found similar results to those described here [12,20,21,14,22,23,18]. However, since these patients did not abstain from PD medication, these results are likely driven by PD treatment rather than PD pathology. This highlights a challenge facing these type of biomarker studies and underscores the need for longitudinal studies and creative ways to correct for treatment effects.

### Peripheral markers of dopamine dynamics

Our analysis identified elevated levels of DOPAL in the plasma of PD patients compared to controls. DOPAL is a cytosolic product of dopamine that is metabolized by monoamine oxidase-A in the outer mitochondrial membrane [24]. DOPAL is a neurotoxin which selectively kills dopamine neurons [25–27]. Toxic DOPAL buildup due to insufficient sequestration by vesicular proteins has been hypothesized to be the precursor to dopamine cell death in PD pathogenesis [28,29]. It is possible that L-dopa treatment in our PD patient population mediated the elevated DOPAL content. However, while in the total patient population L-dopa dosage and DOPAL plasma concentration were associated (r^2^ = 0.365), this correlation was absent when we restricted analysis to PD patients only. We were surprised to observe that DOPAL content differed between PD and control patients in the peripheral blood. The detection of DOPAL in plasma suggests that a test could be devised to measure the DOPAL:DA ratio as an indicator of vesicular dopamine function.

### Differences between PD and control plasma

Despite the confounding effect of L-dopa in this analysis, variations in harmalol and ganglioside GM2 (a metabolite of the glycosphingolipid pathway) associated with PD suggests additional biological changes were detected. Both were identified as elevated in control patient plasma compared to PD patients. To our knowledge, there is not a readily apparent biological mechanism through which PD medication might lower endogenous levels of either harmalol or ganglioside GM2. Therefore, these differences may be due to pathological or physiological differences between patients with and without PD diagnosis, rather than due to an effect of taking PD medication.

Harmalol is an indole beta-carboline alkaloid found in a variety of tissues, including brain, and in a variety of consumable plants, most notably in the leaves of tobacco plants [30–32]. Despite early therapeutic promise [33], studies demonstrating neurotoxic effects of beta-carbolines dampened clinical enthusiasm [34–36]. Later studies found that at low doses, beta-carbolines can be neuroprotective, as demonstrated in PC12 cells [37–39], primary cell culture [40], and in a rat model of PD [41]. Beta-carbolines, particularly harmalol and harmine, exert this effect at the surface of the mitochondrial membrane of dopaminergic cells, preventing oxidative damage from free-radicals [42].

Certainly, our findings that harmalol is elevated in plasma from control patients is exciting given the therapeutic potential of the alkaloid. It is unclear whether control patients display elevated concentrations of harmalol due to an elevated endogenous production of the chemical, or if their elevated harmalol levels are due to a history of consuming of harmalol-containing products. This difference could point to whether individuals who do not develop PD have an antioxidant-mediated protection factor, or whether they engage in habits that lower the risk of PD development. As beta-carboline alkaloids are found in tobacco plants and nicotine is known to independently reduce risk of developing PD [43], the elevated harmalol in control patients may be an artifact of the protective effect of smoking [44]. Although we cannot rule out the possibility, we do not believe it is likely that smoking drove the difference in harmalol in these patients. Concentration of plasma harmalol was not associated with plasma cotinine, a marker for smoking activity (r^2^ = 0.073). Nonetheless, this does not rule out confounding entirely. For instance, participants in the study did not fast prior to serum collection, so it is plausible that dietary differences between cases and controls could account for some of the metabolic variability. Furthermore, while we have no knowledge of link between administration of L-dopa and changes in beta-carboline metabolism, this experiment cannot rule out the effect of PD drugs. In our sample of PD patients, L-dopa plasma concentration did weakly correlate with harmalol concentration (r^2^ = 0.1901), while other studies have shown no correlation [45]. In addition, patients may or may not have been on some other medication that could have confounded the observed metabolic differences. Lastly, it is unknown whether blood plasma harmalol concentration correlates with harmalol levels in the brain. Elevated plasma levels of a compound do not always correspond with elevated levels in the brain [46], and there is evidence that PD patients actually show elevated beta-carboline levels in cerebrospinal fluid [45,47].

Gangliosides are a group of glycosphingolipid compounds that are especially common in the brain but are also found in blood. Gangliosides have been implicated in a variety of neurodegenerative diseases [48]. We identified the ganglioside GM2 and the glycosphingolipid biosynthesis metabolic pathway in our pathway analysis. GM2 levels were lower in PD patients and correlated negatively with motor function as measured by UPDRS-III (r = −0.492). Our work implies that reduced peripheral GM2 could indicate central nervous system dysfunction. GM1 ganglioside is a potential therapeutic target for PD, and a recent study has further suggested GM1’s disease-modifying effect on PD [49,50]. While GM1 has been extensively implicated in human PD, the role of GM2 remains unclear. Other researchers have found that deletion of GM2/GD2 synthase—the enzyme that produces GM2 and GD2 gangliosides—causes a Parkinsonian phenotype, likely due to downstream effects on GM1 [51]. Furthermore, the pathological fibrillation of α-synuclein, which is characteristic of PD, can be inhibited by GM2 (and even more so by GM1) [52]. As with harmalol, confounding—whether due to diet, L-DOPA administration, or some other factor—cannot be ruled out entirely. However, considered together, this analysis points towards ganglioside metabolism and GM2 in particular as potential biomarkers or therapeutic targets for future research.

### Conclusions

Our analysis shows that the presence of L-dopa widely influences the blood plasma metabolome in patients with PD. The influence of L-dopa may be more far-reaching than previously thought: our analysis shows that the effects of PD medications on the metabolome cannot simply be controlled for by removing direct metabolites of the medications. We caution other researchers to seek creative solutions when searching for potential diagnostic and/or prognostic biomarkers in future studies. Lastly, we highlight two metabolites of interest that were elevated in control versus PD serum, harmalol and the ganglioside metabolic pathway.

## Materials and Methods

This study was approved by the human subject committee of Emory University. Subjects participated after written informed consent was obtained in accordance with the Declaration of Helsinki.

### Study Population Characteristics

The plasma samples analyzed were collected from patients recruited through the Emory Movement Disorders Clinic and controls recruited in the Atlanta area from 2012–2013. The final study population contained 21 PD patients and 13 control patients. Demographic information was collected from all participants and is summarized in Table 1.

### Clinical Data Collection

Clinical data was collected from all PD patients, including disease duration and daily L-dopa equivalent dosage of antiparkinsonian medications. All participants had UPDRS-III motor assessments by a fellowship-trained movement disorders neurologist [53]. All participants had cognitive assessments with the Montreal Cognitive Assessment (MOCA), and all participants completed the Non-Motor Symptoms Questionnaire (NMSQ) [54,55]. Prior to plasma collection, patients were not required to fast nor to stop their PD medications. Exams and sample collections were typically conducted between 9 am and 11 am. Blood samples were collected in EDTA tubes, and immediately placed on ice. They were then transferred on ice an centrifuged at 2200 RPM for 15 min. The plasma was transferred to a transport vial and frozen at −80*C until mass spectrometry analysis. The time from sample collection to storage in the −80*C freezer was less than 2 hours.

### Mass spectrometry

Samples were prepared as previously described [12]. Briefly, 65 μL of plasma was treated with 130 μL of acetronitrile containing a mixture of 14 stable isotope standards. After mixing and incubation at 4°C for 30 min, precipitated proteins were pelleted via centrifugation for 10 min at 16,100 x *g* at 4°C. Supernatants were transferred to autosampler vials and stored at 4°C until analysis. Sample extracts were analyzed in triplicate by liquid chromatography-Fourier transform mass spectrometry (Dione Ultimate 3000;Thermo Scientific Q-Exactive HF High-Resolution Mass Spectrometer) with 10 uL injection volume and a formic acid/acetonitrile gradient as described previously [56]. Electrospray ionization was used in the positive ion mode. Mass spectral data were collected at a resolution of 70,000 and scan range 85 to 1250 *m/z* [11,57]. Raw data files were extracted and aligned using apl CMS with modifications by xMSanalyzer with each feature uniquely defined by *m/z* (mass-to-charge ratio), retention time, and sample ion intensity (integrated ion intensity for the peak).

### Metabolomics data analysis

R was used to analyze the metabolomics data. We first eliminated features which had greater than 35% variability within technical replicates. This reduced the data set from a total of 10,471 m/z values to 9,071 m/z values. Features detected only in PD cases were not removed from the dataset, as the potential measurement of exogenous compounds was one of the major aims of this study. Since a detection value of zero did not imply the absence of the feature, but rather that the amount of feature present was below the threshold of detection, we imputed half of the minimum detected value to all zero values. We then applied a generalized log transformation as described in MetaboAnalyst (55), which resulted in an approximately normal distribution.

To determine which features differed between PD versus control plasma, two-sample t-tests were conducted for each feature. P-values were adjusted using the FDR correction; features were sorted using the adjusted p-values. This analysis was performed using MetaboAnalyst software [58] and original R code to verify results. To determine which metabolic pathways were most affected by PD drug treatments, we also performed a metabolome-wide association study restricted to PD patients wherein we calculated the correlations of the abundance of primary PD drug metabolite *(m/z* = 212.092, retention time = 71) with every other metabolite feature, then created a network of metabolites that most significantly correlated with PD.

The degree of association between two variables was analyzed via linear correlation analysis and reporting the r squared value and direction of association where appropriate.

Pathway analyses were performed in Mummichog [59]. The “force primary ion” option was chosen, ensuring that any predicted metabolites were present in at least their primary adduct (M+H^+^). A *p* threshold of 0.05 was selected. Pathways identified in Mummichog were visualized in Cytoscape [60]. Identity of networks was visualized using Tableau (Seattle, Washington).

For orthogonal partial least squares-discriminant analysis (OPLS-DA), the data was mean-centered prior to analysis [61,62]. OPLS-DA was performed and visualized both using R and MetaboAnalyst tools.

Putative identities of metabolite features was determined via multiple parallel methods: a custom deconvolution and identification algorithm (Emory University, Atlanta, Georgia), xMSanalyzer [57], and manual lookup in various online metabolite databases (Human Metabolome Database, Kyoto Encyclopedia of Genes and Genome) [63,64].

---

**Supplementary Table 1.**
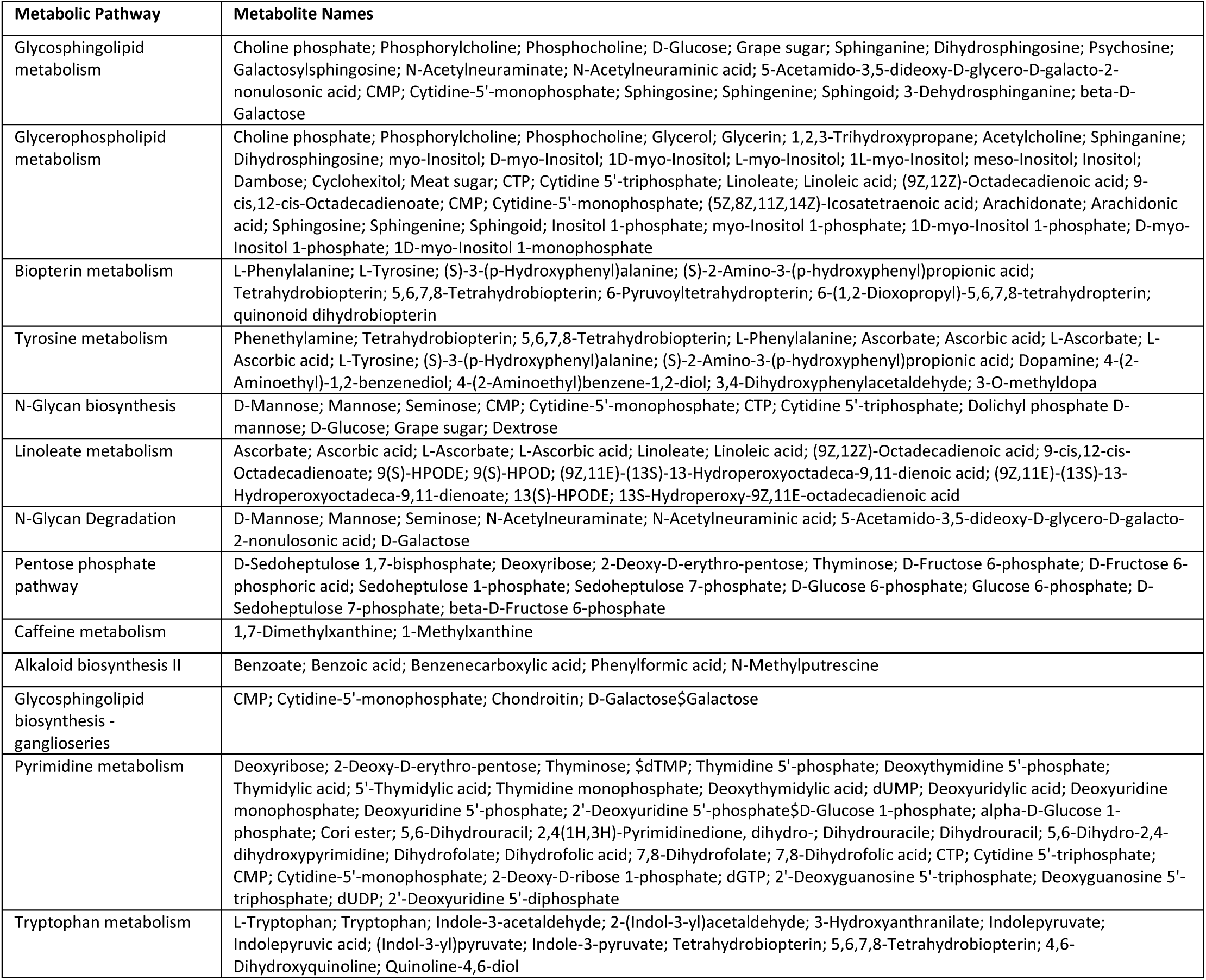
13 metabolic pathways identified as significantly different between PD patients and control patients. Mummichog software identified metabolic pathways that are most likely to be different between the PD and control patients. Identities of compounds found within these pathways are also listed.

**Supplementary Table 2.**
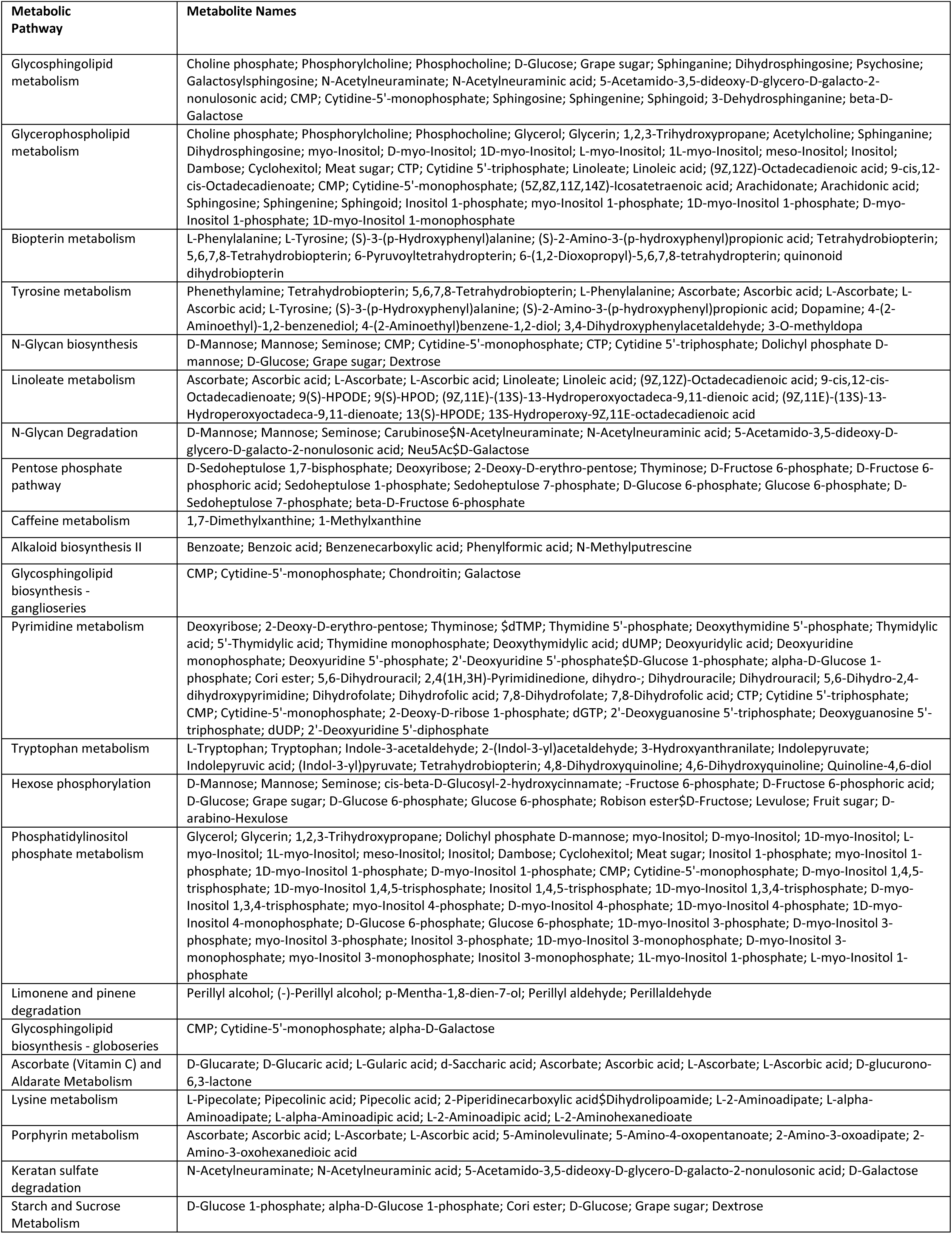
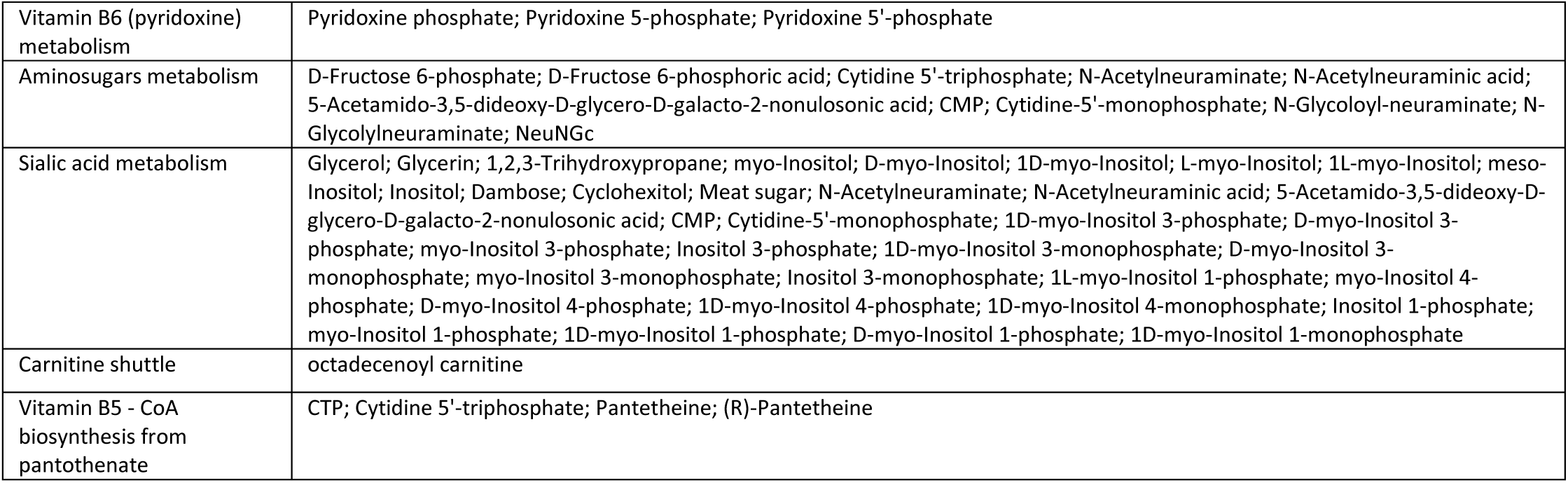
27 metabolic pathways identified as correlated with L-DOPA. Mummichog software identified metabolic pathways that are strongly associated with the presence of L-DOPA. Identities of compounds found within these pathways are also listed.

**Supplementary Fig 1.**
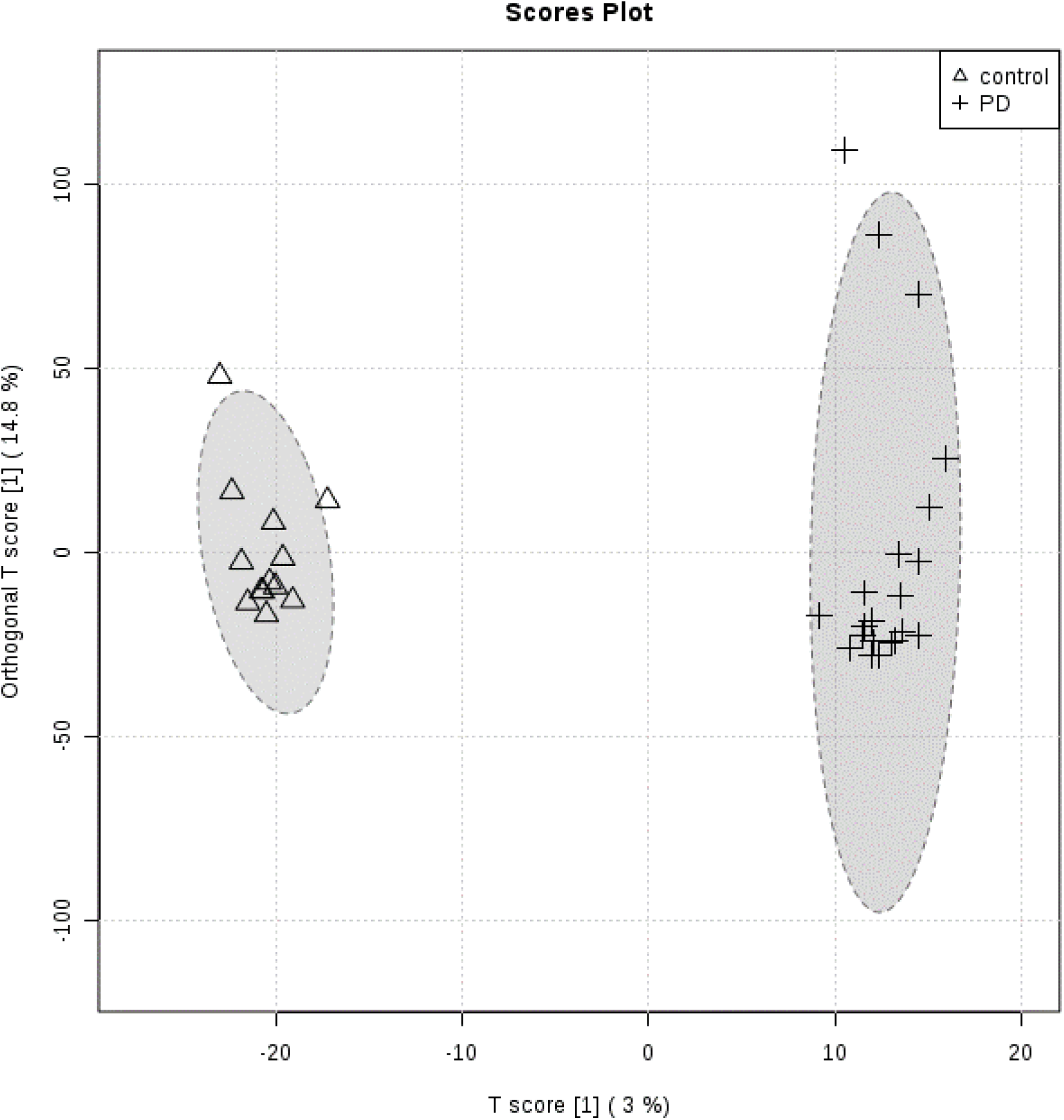
OPLS-DA between PD and control metabolomes with PD drug metabolites manually removed.

**Supplementary Fig 2.**
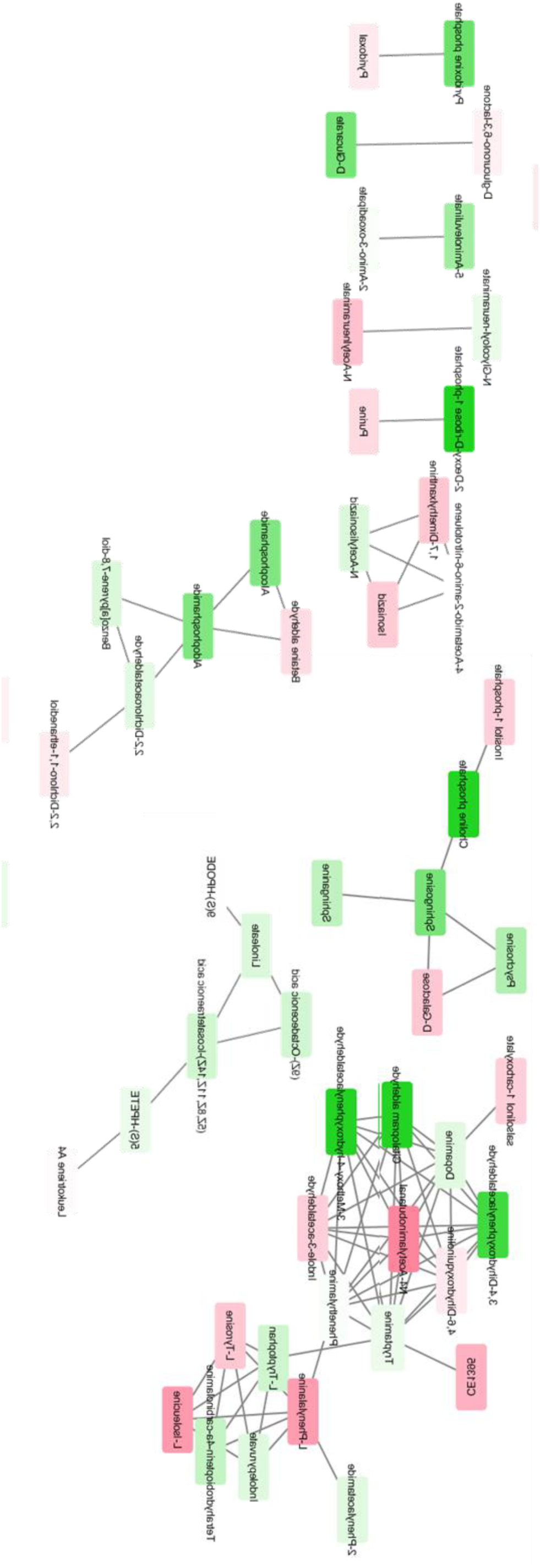
(rotated 90 degrees for legibility) Network analysis of PD plasma analytes that correlate with l-dopa. Spearman correlation between each *m/z* and L-dopa was calculated among PD patients. Darker shades of green correspond to stronger positive Spearman correlations; darker shades of red correspond to stronger negative Spearman correlations.

**Supplementary Fig 3.**
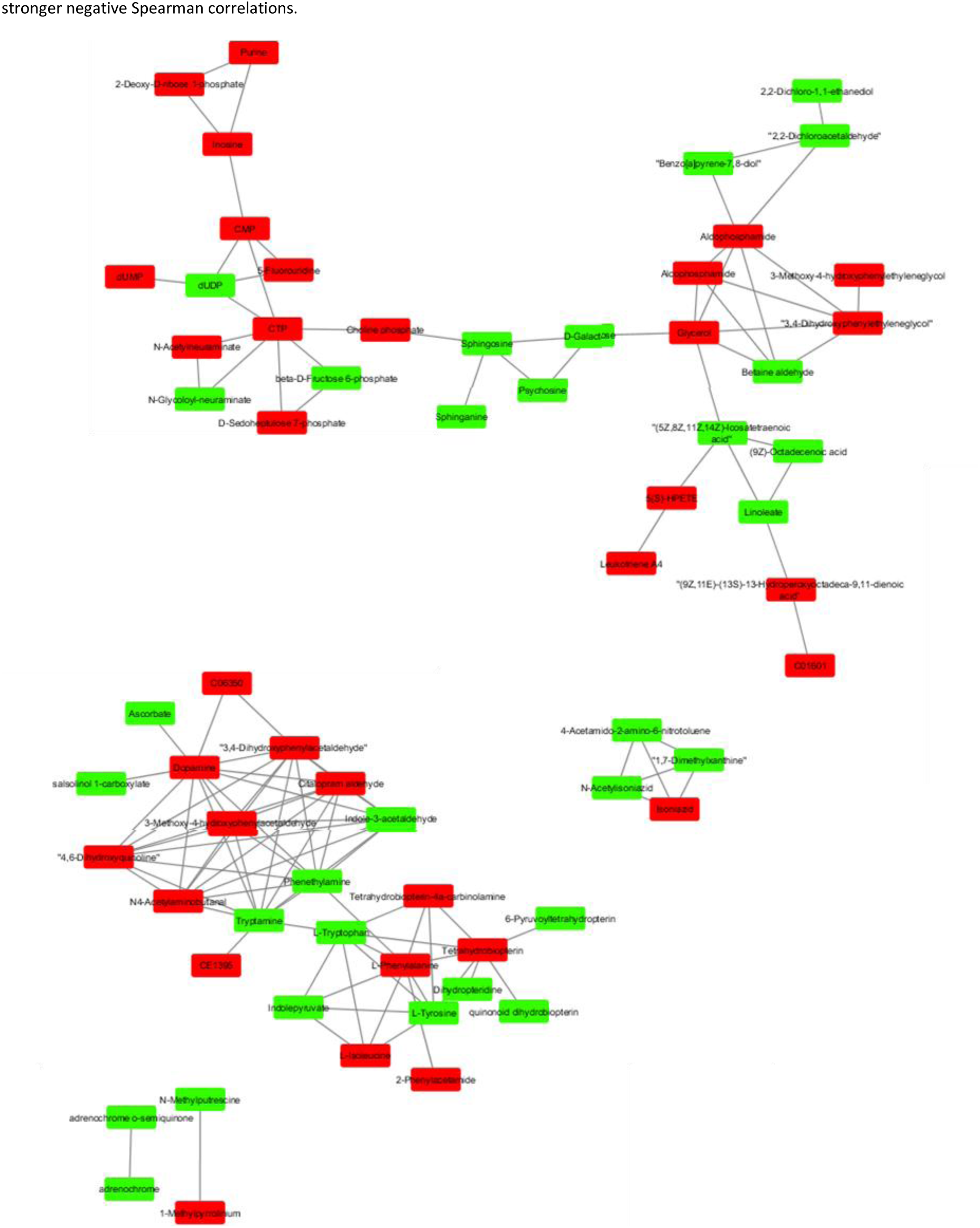
Network analysis of the metabolites that are most different between PD and control patients. Red nodes correspond to metabolites elevated in PD patients compared to controls, while green nodes correspond to metabolites lower in PD patients compared to controls.

